# Aligning Distant Sequences to Graphs using Long Seed Sketches

**DOI:** 10.1101/2022.10.26.513890

**Authors:** Amir Joudaki, Alexandru Meterez, Harun Mustafa, Ragnar Groot Koerkamp, André Kahles, Gunnar Rätsch

**Affiliations:** Department of Computer Science, ETH Zurich, Zurich 8092, Switzerland; University Hospital Zurich, Biomedical Informatics Research, Zurich 8091, Switzerland; Swiss Institute of Bioinformatics, Lausanne 1015, Switzerland; ETH AI Center, 8092 Zurich, Switzerland

**Author notes:** Equal contribution (Amir Joudaki, Alexandru Meterez).

**Keywords:** sequence-to-graph alignment, tensor sketching, tensor embedding

## Abstract

Sequence-to-graph alignment is an important step in applications such as variant genotyping, read error correction and genome assembly. When a query sequence requires a substantial number of edits to align, approximate alignment tools that follow the seed-and-extend approach require shorter seeds to get any matches. However, in large graphs with high variation, relying on a shorter seed length leads to an exponential increase in spurious matches. We propose a novel seeding approach relying on long inexact matches instead of short exact matches. We demonstrate experimentally that our approach achieves a better time-accuracy trade-off in settings with up to a 25% mutation rate.

We achieve this by sketching a subset of graph nodes and storing them in a *K*-nearest neighbor index. While sketches are more robust to indels, finding the nearest neighbor of a sketch in a high-dimensional space is more computationally challenging than finding exact seeds. We demonstrate that if we store sketch vectors in a *K*-nearest neighbor index, we can circumvent the curse of dimensionality. Our long sketch-based seed scheme contrasts existing approaches and highlights the important role that tensor sketching can play in bioinformatics applications. Our proposed seeding method and implementation have several advantages: i) We empirically show that our method is efficient and scales to graphs with 1 billion nodes, with time and memory requirements for preprocessing growing linearly with graph size and query time growing quasi-logarithmically with query length. ii) For queries with an edit distance of 25% relative to their length, on the 1 billion node graph, longer sketch-based seeds yield a 4× increase in recall compared to exact seeds. iii) Conceptually, our seeder can be incorporated into other aligners, proposing a novel direction for sequence-to-graph alignment.

The implementation is available at: https://github.com/ratschlab/tensor-sketch-alignment.

## 1 Introduction

Our work focuses on *sequence-to-graph alignment*, defined as aligning a query sequence to a sequence graph [37,15]. Sequence-to-graph alignment involves finding the minimal number of *editing operations* to transform the query to a reference sequence encoded in the graph. While there are various cost schemes for penalizing edits, edit distance assigns a cost of 1 to *substitution, insertion*, and *deletion* on the query.

Since the time complexity of optimal sequence-to-graph alignment grows linearly with the number of edges in the graph [20,16], many approaches instead follow an approximate *seed-and-extend* strategy [2], which operates in four main steps: i) *seed extraction*, which in its simplest form involves finding all substrings with a certain length, ii) *seed anchoring*, finding matching nodes in the graph, iii) *seed filtration*, often involving clustering [9,37] or co-linear chaining [25,1,32,8] of seeds, and iv) *seed extension*, involving performing semi-global pairwise sequence alignment forwards and backwards from each anchored seed [28]. We will review the usage of exact seeds utilized in tools such as vg[15] and GraphAligner [37] and discuss their limitations in a high mutation-rate setting.

Let us highlight the limitations of *k-mers*, defined as substrings with length *k*, as candidates for seeding, with an example.

### Example 1.

For reference sequence *S* ∼ {`A,C,G,T}`^*N*^, for *i* = 1 up to 100, with probability 1 − *r* copy the *i*-th character *q*_*i*_ ← *S*_*i*_, and with probability *r* mutate it *q*_*i*_ ∼ *∑* \ {*S*_*i*_}. The expected number of hits i.e. common *k*-mers between query and reference, is at most (100/*k*)(1 − *r*)^*k*^. On the other hand, the expected number of spurious hits is *N* 4^−*k*^.

The simple setting in Example 1 illustrates the challenge many alignment methods face in practice. Since using a large *k* for seed length reduces the number of spurious matches, methods such as BWA-MEM [28], E-MEM [26], deBGA [31], and PuffAligner [1] find maximal exact matches (MEMs) between the read and the reference genome. However, the higher precision comes at the expense of fewer seeds and lower recall. For example, for *k* = 21 in Example 1, finding an exact seed for a query with mutation rate *r* = 0.25 becomes exceedingly rare, the expected number of hits being (100/21)(1 − 0.25)^21^ ≈ 0.01.

The alternative is to use shorter seeds to increase the likelihood of finding accurate alignments. Methods that use De Bruijn graphs as graph model [5], or an auxiliary index [40], are restricted to finding seeds of minimum length *k*. To find seeds when the query has a high edit distance relative to the reference sequences, sequence-to-graph alignment tools will either set *k* to a small value [30,15,37] or use a variable-order DBG model [4,3,24,25] to also allow for anchoring seeds of length less than *k* [24,25]. However, shorter seeds generally lead to a more complex and connected graph topology which can lead to a combinatorial explosion of the search space. In the same setting as Example 1, with *N* = 10^9^ and *k* = 12, there will be 10^9^/4^12^ 60 false positive matches for every true positive match, while for *r* = 0.25 mutation rate the recall will be (100/12)(1 − 0.25)^12^ ≈ 0.26. Therefore, any attempt of changing *k* will either risk a lower recall or a higher false positive.

Due to the inherent shortcomings of short and long exact matches, Břinda, et al. [6] propose *spaced seeds* to incorporate long seeds at higher mutation rates by masking out some positions in the seed. For example, using 0101 as a mask, “A<u>C</u>G<u>T</u>” will match with “G<u>C</u>A<u>T</u>”. While spaced seeds were shown to be more sensitive than contiguous seeds without increasing the number of spurious matches, they assume that only mismatches occur in the alignment with no *insertions* or *deletions (indels)* [33].

In this work, we propose using long inexact seeds based on Tensor Sketching [23]. We use a succinct *De Bruijn graph (DBG)* [5] as the graph model, and preprocess it in two main stages: i) A subset of nodes in the DBG is sketched into a vector space, while insertions, deletions, and substitutions are approximately embedded into the *L*^2^ metric. ii) To be able to efficiently retrieve similar sketch vectors, the sketches of nodes are stored in a *Hierarchical Navigable Small Worlds (HNSW)* [34] index. Aligning a query sequence involves three main stages: i) Compute sketch vectors for all *k*-mers in the query. ii) Use the HNSW index to find the nearest vectors as candidate seeds. iii) Extend these seeds backwards and forwards to find the best possible alignment. Figure 1 gives an overview of our sketch-based scheme.

**Fig. 1:**
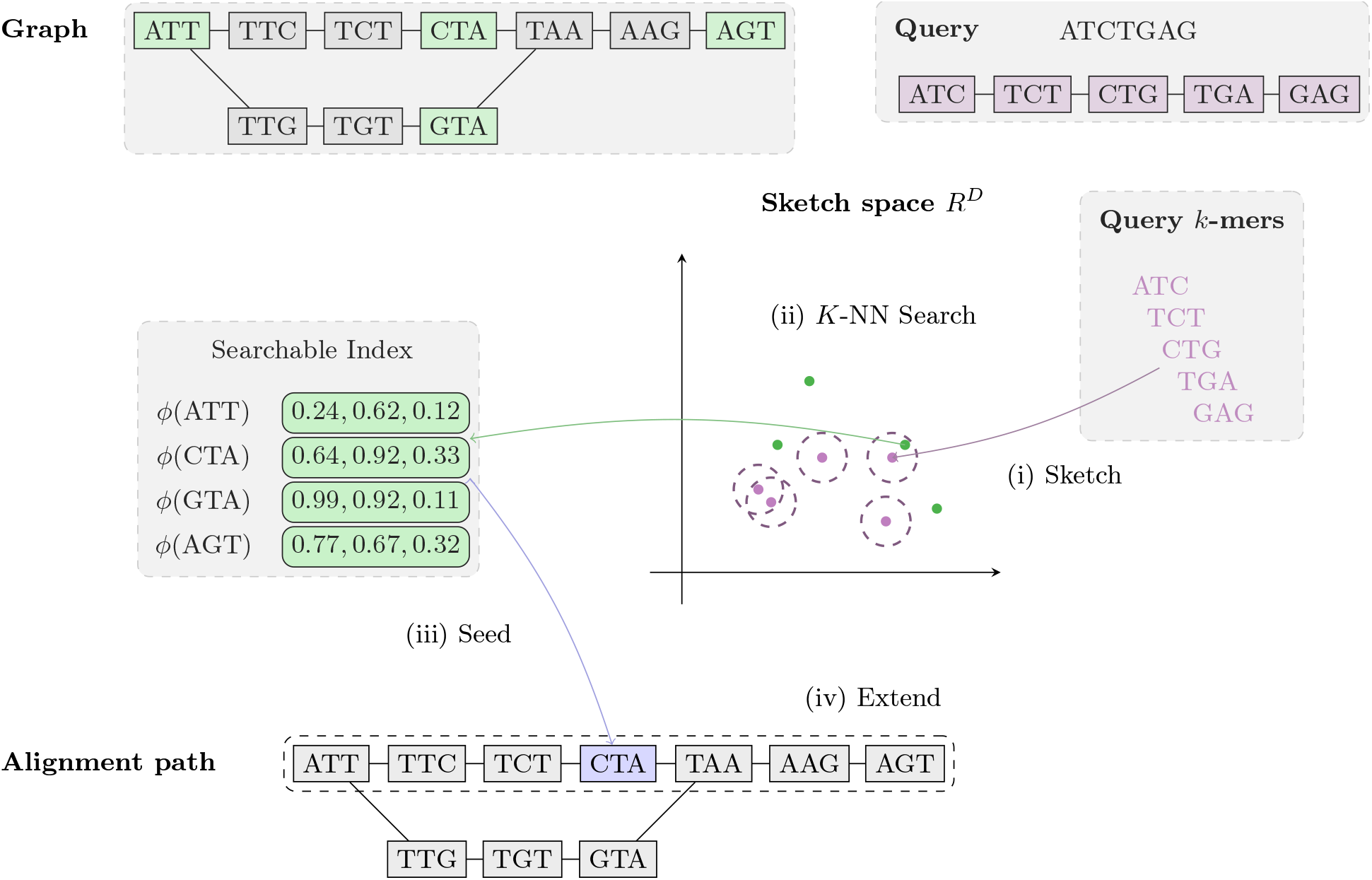
The full sequence-to-graph alignment of one query sequence using MG-Sketch consists of several steps: i) Sketching the spell of every graph node in the *k*mer-cover of the DBG and storing the sketches in a hierarchical searchable index, mapping from sketch vector to node, ii) For every *k*-mer in the query (magenta), find the nearest neighbors with the smallest sketch *L*^2^-distance (green) stored in the index, iii) Seed an alignment at every node returned in the previous step and iv) Extend forwards and backwards from each seed, finding the path in the graph that maximizes the alignment score of the query.

### Structure of the manuscript

In Section 2.1, we introduce our notation, formalize the problem of sequence-to-graph-alignment, and explain how tensor sketching estimates edit distances. In Section 2.3, we introduce the hierarchical index for inexact search, and in Section 2.4, we empirically compare our method against other tools. Section 3 is dedicated to our experimental validations, starting with the synthetic sequence generation in Section 3.1. The accuracy and scalability of our method are covered in Sections 3.3 and 3.2, respectively. Finally, we present the limitations of sketch-based seeds, as well as future directions in Section 4.

## 2 Methods

### 2.1 Preliminaries

#### Terminology and notation

We use [*N*] := {1, …, *N*}. *∑* denotes the alphabet, e.g., the nucleotides {`A,C,G,T}`, or amino acids. For *k* ∈ ℕ^+^, *∑*^⋆^ denotes the set of all strings over *∑*, and *∑*^*k*^ denotes the subset of all strings with length *k*. Throughout the text, we use the terms string and sequence interchangeably to refer to members of *∑*^⋆^. *q*[*i*] and *q*_*i*_ denote the *i*-th character of the sequence, and *q*[*i* : *j*] := *q*_*i*_*q*_*i*+1_ … *q*_*j*_ denotes the sliced substring from *i*, up to *j. k*-mers(*s*) denote the substrings of length *k* in string *s*: *k*-mers(*s*) := {*s*[*i* : *i* + *k* − 1] : 1 ≤ *i* ≤ |*s*| − *k* + 1}. *p* ∘ *q* denotes the concatenation of *p* and *q*. The *edit distance* ed(*p, q*), also referred to as the Levenshtein distance [41], is defined as the minimum number of insertion, deletion, and substitution operations needed to transform one sequence into the other. The normalized edit distance, is defined as edit distance divided by maximum length 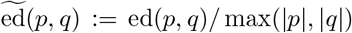. Throughout the manuscript, the term “mutation” refers to the normalized edit distance, serving as an abstraction for the combined biological sources of variation and errors in sequencing.

#### Succinct De Bruijn graph (DBG)

The reference DBG is a directed graph over the reference sequences that encodes all substrings of length *k* as vertices and all substrings of length *k* +1 as directed edges. Formally, let 𝒮 = {*r*_1_,…, *r*_| 𝒮 |_} be the input sequences. The reference graph is a directed graph *G* = (*V, E*), with vertices *V* := ∪_*s* ∈ 𝒮_ *k*-mers(*s*) and edges *E* := {(*u, v, v*[*k*]) ∈ *V* × *V* × *∑* : *u*[2 : *k*] = *v*[1 : *k* − 1]}. The label of edge *e* = (*u, v, v*[*k*]) ∈ *E* is denoted by *l*_*e*_ := *v*[*k*] ∈ *∑*.

#### Sequence-to-graph alignment

Define 𝒲 _*G*_ as the set of all walks, where each walk *w* ∈ 𝒲 _*G*_ is defined as a list of adjacent edges *w* = (*v*_0_, *v*_1_, *l*_1_), …, (*v*_|*w*|−1_, *v*_|*w*|_, *l*_|*w*|_) ∈ *E*^| *w* |^. Define the *spelling* of a walk as the concatenation of the first *k*-mer on the walk, with the labels of the rest of the edges on the same walk *π* (*w*) := *v*_0_ ∘ *l*_1_*l*_2_ … *l*_|*w*|_. Thus, given the query sequence *q* ∈ *∑* ^⋆^ and the reference graph *G*, the optimal *alignment* is the set of walks which achieves the minimum edit distance between the query sequence and the spelling of the walk 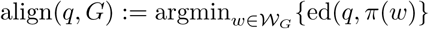.

### 2.2 Estimating edit distance using Tensor Sketch

Our approach to the seed extraction and anchoring steps of seed-and-extend (see Section 1) is based on an index of *k-mer sketches* rather than *k*-mers. Briefly, we compute Tensor Sketches [23] of the graph nodes and store them in a nearest-neighbor search index [22] that maps each sketch to its corresponding graph node.

Tensor Sketching (TS) maps sequences to a vector space that embeds the edit distance into the 𝓁 ^2^ metric. Conceptually, TS can be described in two steps: i) Tensor Embedding (TE), which counts how many times each subsequence appears in the sequence, ii) Implicit tensor sketching, which lowers the dimension without constructing the larger tensor embedding space.

#### Tensor Embedding

Given sequence *a* ∈ *∑* ^*n*^, define ℐ as set of increasing subsequences of length *t*: ℐ := {(*i*_1_,…, *i*_*t*_) ∈ [*n*]^*t*^ : *i*_1_ < … < *i*_*t*_}, and for all *s* ∈ *∑* ^*t*^, define tensor embedding *T*_*a*_[*s*] as the count of all subsequences of length *t* in *s*: *T*_*a*_[*s*] := # {*I* ∈ ℐ: *a*[*i*_1_,. ‥, *i*_*t*_] = *s*}. We can view *T*_*a*_ as a |*∑*|^*t*^-dimensional tensor, with the alphabet as its indices. Given sequences *a, b*, we approximate the edit distance by 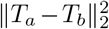 up to a constant factor. Figure 2 illustrates how the embedding distance approximates edit operations.

**Fig. 2:**
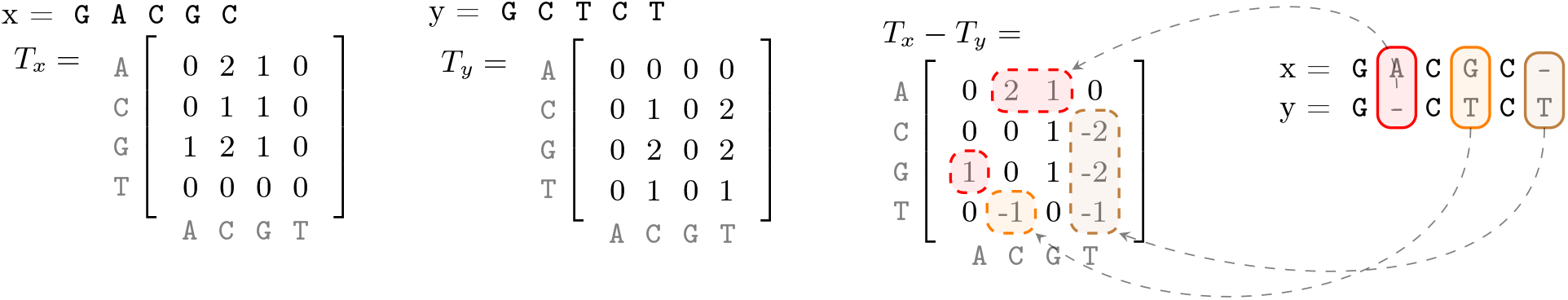
Tensor Embedding illustration for *t* = 2. Observe that the substitution, insertion, and deletion, correspond to blocks of non-zero elements in the difference tensor.

#### Tensor embedding preserves Hamming distance under 𝓁^2^-norm

Intuitively, tensor embeddings are robust to mutations because they count and sketch subsequences instead of *k*-mers. We can define normalized tensor embedding distance *d*_*te*_ between two sequences *a* and *b* as:

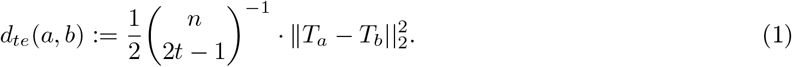

The following lemma demonstrates that tensor embedding preserves edit distance:

##### Lemma 1.

*Let a be a uniform random sequence of length n in ∑* ^*n*^, *and for a fixed mutation rate r* ∈ [0, 1] *let b be a sequence where a*_*i*_ *is substituted by a new character b*_*i*_ ∈ *Unif* (*∑* \*a*_*i*_) *with probability r and b*_*i*_ = *a*_*i*_ *otherwise. Then for n* ≫ 2*tα:*

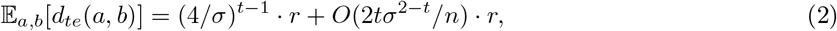

*which for DNA with α* =4 *and fixed t gives* 𝔼 [*d*_*te*_(*a, b*)] = (1 + *O*(*n*^−1^)) · *r*.

Note that the (4/*σ*)^*t*−1^ factor does not depend on the sequences. Therefore, Lemma 1 provides a guarantee that the average distance of mutated pairs remains within a linear estimate of the mutation. Observe that the edit distance in this setting will be *nr*. Therefore, we can, alternatively, refer to *r* as edit distance normalized by length. The proof of Lemma 1 is given in Appendix B. For numerical validation of this bound, see Figure 3.

**Fig. 3:**
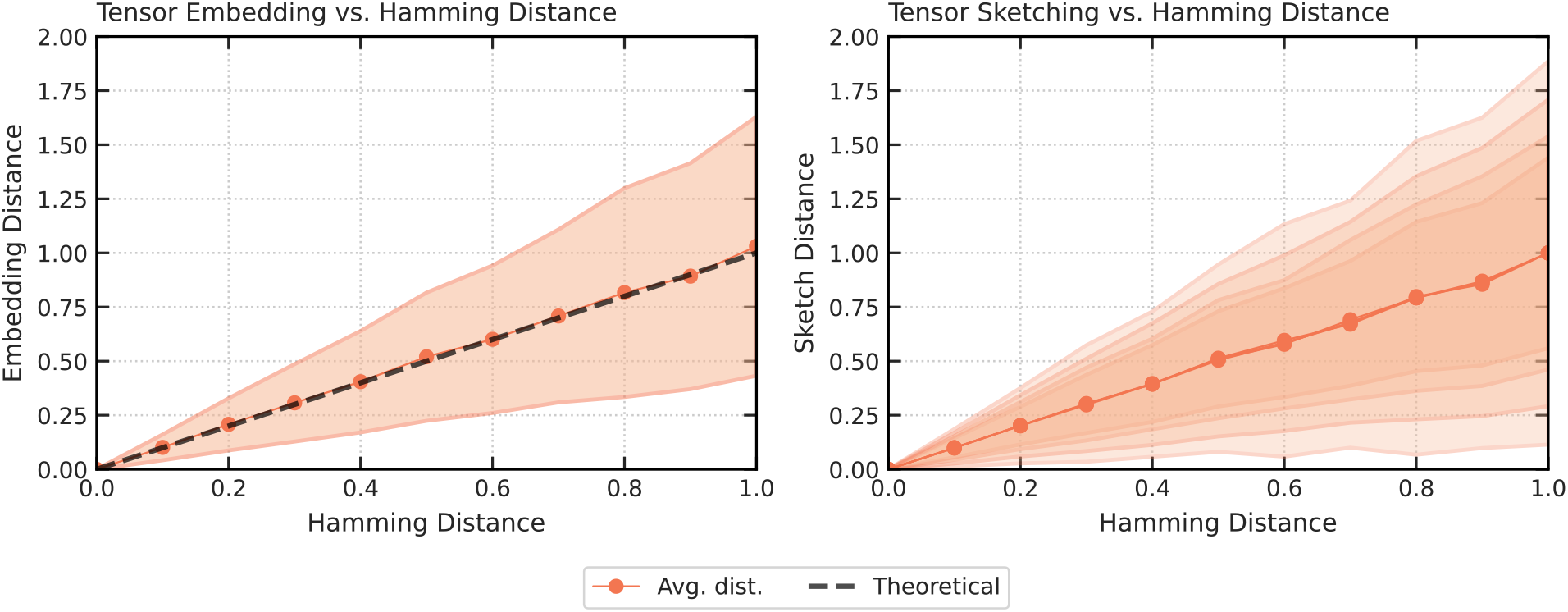
(left) Tensor Embedding vs. Hamming Distance: Dashed line represents the closed-form expectation according to the Lemma 1 with *t* = 3, *n* = 1000, which is almost overlapping with the empirical average represented by the solid line. The solid line shows the mean tensor embedding distance with *t* = 3, averaged over 2000 sequence pairs of length 1000, at normalized Hamming distances ranging from 0 to 1. (right) Tensor Sketching vs. Hamming Distance: Average tensor sketch distance on the same dataset used for tensor embedding distances, with the solid line showing the average sketch distance with *t* =3 and *D* = 8, 16, 32, 64. The sketch distance is normalized such that it is equal to 1 for Hamming distance 1. Shaded regions represent the standard deviation of the sketch distance for *D* = 8, 16, 32, 64, with the lightest shade corresponding to *D* = 8. (right).

#### Tensor Sketching

Tensor Sketching is an implicit, Euclidean-norm preserving dimensionality reduction scheme. Crucially, it projects |*∑*|^*t*^ -dimensional tensors onto constant *D* ∈ ℕ ^+^ dimensions and linearly preserves 𝓁 ^2^ distances, without ever constructing the tensors. We define the tensor sketching function 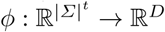, that embeds an |*∑*|^*t*^-dimensional tensor into ℝ^*D*^. Given pairwise independent hash functions *s*_*i*_, *h*_*i*_ : *∑* → [*D*] and *s*_*i*_ : *∑* → {−1, 1}, define the tuple hash 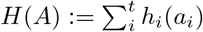, and tuple sign hash 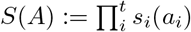, where *A* = (*a*_*i*_)_*i*≤ *t*_ ∈ *∑* ^*t*^. Finally, the tensor sketch *ϕ* for a tensor 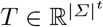 is defined as 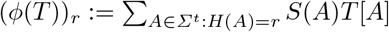 for all *r* ∈ [*D*].

Crucially, we have 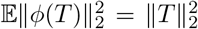 and bounded second moments 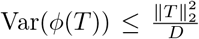 (See Lemma 7 of [36]). Figure 3 shows lower and upper bounds. Finally, [23] reports that *Tensor Slide Sketch (TSS)*, which concatenates tensor sketches of windows *w*, using a stride *Δ*. within a given *k*-mer, improves the sensitivity to smaller edit distances. Therefore, we used TSS for our seeding scheme (See Appendix C for more details).

### 2.3 Anchoring seeds with a hierarchical search index

The final pre-processing step is to build a *K*-Nearest Neighbor index of sketches. Conceptually, we can represent the search index as a function from vector space ℝ^*D*^ to a list of graph nodes *K*-NN(*v*)= (*v*_*1*_,…, *v*_*K*_), for some pre-determined number of neighbors *K* ∈ ℕ^+^. During the alignment, we anchor every seed in the query *s* ∈ *k*-mers(*q*), to the vertices returned by the index *K*-NN(*ϕ* (*s*)).

Note that if *v* and *u* are within *s* steps on the graph, they share 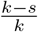 of their sequence content. To avoid indexing redundant information, we only store a *k*cover of the graph, defined as a subset of vertices such that any walk on the graph with over *k* nodes contains at least one node in the *k*cover. For any walk *W* ∈ 𝒲 _*G*_, | *W* | *≥ k*, it holds that *W* ∩ *k*cover(*G*) ≠ Ø. Observe that if the optimal alignment walk for a query has over *k* nodes, it suffices to sketch the *k*-cover to anchor at least one of its seeds.

While sketch vectors are more robust to mutations than exact seeds, retrieving nearest neighbors in a high dimensional space faces the *curse of dimensionality* [27]. *Locality Sensitive Hashing [19,10]* is the first method to get a constant approximation for the nearest neighbors problem, with a theoretically proven sub-linear time with respect to dataset size. However, to scale to billions of nodes in the genome graphs, it is crucial to store sketch vectors in a *Hierarchical Navigable Small Worlds (*HNSW) index [34]. We use the efficient implementation of HNSW from Facebook AI Similarity Search (FAISS) [22]. The pseudo-code for the anchoring is presented in Algorithm 1.

#### Algorithm 1: Anchoring

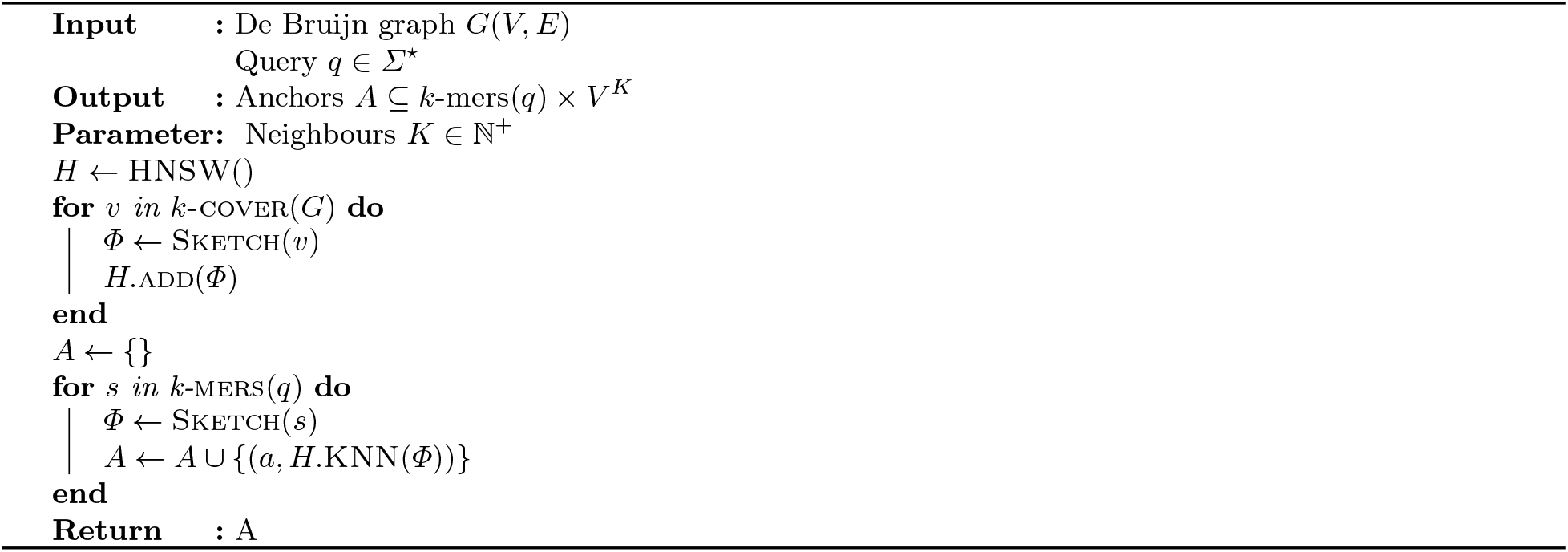

### 2.4 Adjusting extension for misaligned anchors

MetaGraph Align (MG-Align) follows a seed-and-extend approach, with a dynamic program to determine which path to take in the graph, producing a semi-global alignment. We made a few modifications to adjust for misaligned anchors in the MG-Sketch seeder. To demonstrate this issue, let *v*_1_, *v*_2_,…, *v*_*M*_ denote a list of adjacent *k*-mers on the graph, and let *s*_1_,…, *s*_|*q*|-*k*+1_ denote seeds in the query *s*_*i*_ := *q*[*i* : *i* + *k* − 1]. Let us assume that if *s*_*i*_ is anchored to *v*_*j*_, the alignment will be optimal. Observe that ed(*s*_*i*_, *s*_*i*+*δ*_) ≤2*δ*, obtained by deleting the initial *δ* characters from *s*_*i*_ and inserting the last *δ* characters of *v*_*i*+*δ*_. By the triangle inequality, if ed(*s*_*i*_, *v*_*j*_) ≤ *d*, it holds that ed(*s*_*i*+_, *v*_*j*_) ≤*d* + 2*δ*. While Tensor Sketching preserves the edit distance, due to inherent stochasticity in sketching and retrieval, *s*_*i*+*δ*_ may be anchored to *v*_*j*_, instead of *s*_*i*_ to *v*_*j*_. This may produce an additional cost of 2*δ* during the forward and backward extension. To avoid this unnecessary cost, during the forward extension, indels occurring in the beginning of the query are free. If the forward extension completes, we initiate the backward extension from the position of the first matching positions.

## 3 Experimental results

We implemented the MetaGraph Sketch (MG-Sketch) algorithm, which uses our novel sketch-based seeder, in the MetaGraph [24] framework. We primarily compare against MetaGraph Align (MG-Align), which is also implemented in MetaGraph, but uses an exact seeding scheme. We compare MG-Sketch against GraphAligner (GA) [37], vg map [18] and vg mpmap [39] on both synthetic and real datasets (See Appendix A for more details).

### 3.1 Synthetic Data Generation

We generate a synthetic dataset by initializing the sequence pool with *𝒮*_0_ {*s*_0_}, where *s*_0_ is a random sequence *s*_0_ ∼ *∑* ^*N*^, for some fixed length *N* ∈ ℕ^+^. At each level *i* ∈ ℕ^+^, we randomly mutate all sequences in the pool by independently substituting every character with 1% probability. We add the mutated sequences to the pool 𝒮 _*i*_ ← ∪ 𝒮_*i*−1_ ⋃ {mutate(*s*): *s* ∈ 𝒮_*i*−1_}, doubling the pool size. We repeat this process up to ℓ levels.

We build a De Bruijn graph *G* using MetaGraph [24], on the synthetic sequences 𝒮 _𝓁_ with *N* = 1000 and *k* = 80. Recall that all sequences in *S*_𝓁_ are of size *N*, resulting in 2^𝓁^*N* nodes in the graph. Construction of *S*_𝓁_ ensures that approximately 1% of the nodes in *G* are “branching”, i.e. they have at least one incoming or outgoing connection, thereby increasing the difficulty of the sequence-to-graph alignment.

To generate ground truth sequences for alignment, we start with a random walk *w*_*i*_ in the graph with 250 edges, yielding reference sequences *s*_*i*_ := *π*(*w*_*i*_), with length |*π*(*w*_*i*_)| = 80 + 250 = 330. To obtain the query, we apply mutations with rates 5%, 10%, 15%, 20%, and 25% to the references *q*_*i*_ := mutate_*r*_(*s*_*i*_). Conceptually, the alignment method *f* takes the query *q*_*i*_ and graph *G* as input, and returns a candidate spell *f*(*q*_*i*_, *G*) as output. We quantify the optimality of *f* by measuring the edit distance between the reference and the candidate spell: ed (*s*_*i*_, *f*(*q*_*i*_)). We define recall as ed (*s*_*i*_, *f*(*q*_*i*_)) ≤ *k*λ, where λ ∈[0, 1] controls the accuracy of the alignment (lower is more accurate). For all experiments, we set λ = 0.1. Finally, we average the recall and alignment time per query over 500 samples at each mutation rate:Recall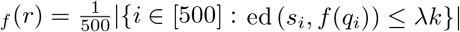.

### 3.2 Sketch-based alignment achieves high recall in high mutation settings

We show that MG-Sketch achieves a uniformly higher recall than MG-Align. Figure 5 shows that MG-Sketch outperforms MG-Align across all graph sizes, particularly in high mutation regimes. MG-Sketch reaches 3 × and 1.8× higher recall than MG-Align on 25% and 20% mutation rates respectively, while getting a comparable or better recall in all other cases. If MG-Align does not find exact 80-mer matches, it falls back to shorter seeds, until a minimum seed length of 15. If we compute the expected number of hits analogous to Example 1, we get (330/15)(1 − 0.25)^15^ ≈ 0.29. This is comparable to the empirical value for 25% mutation in Figure 5. Remarkably, MG-Sketch exceeds this recall by a significant margin, highlighting the importance of long inexact seeds.

**Fig. 4:**
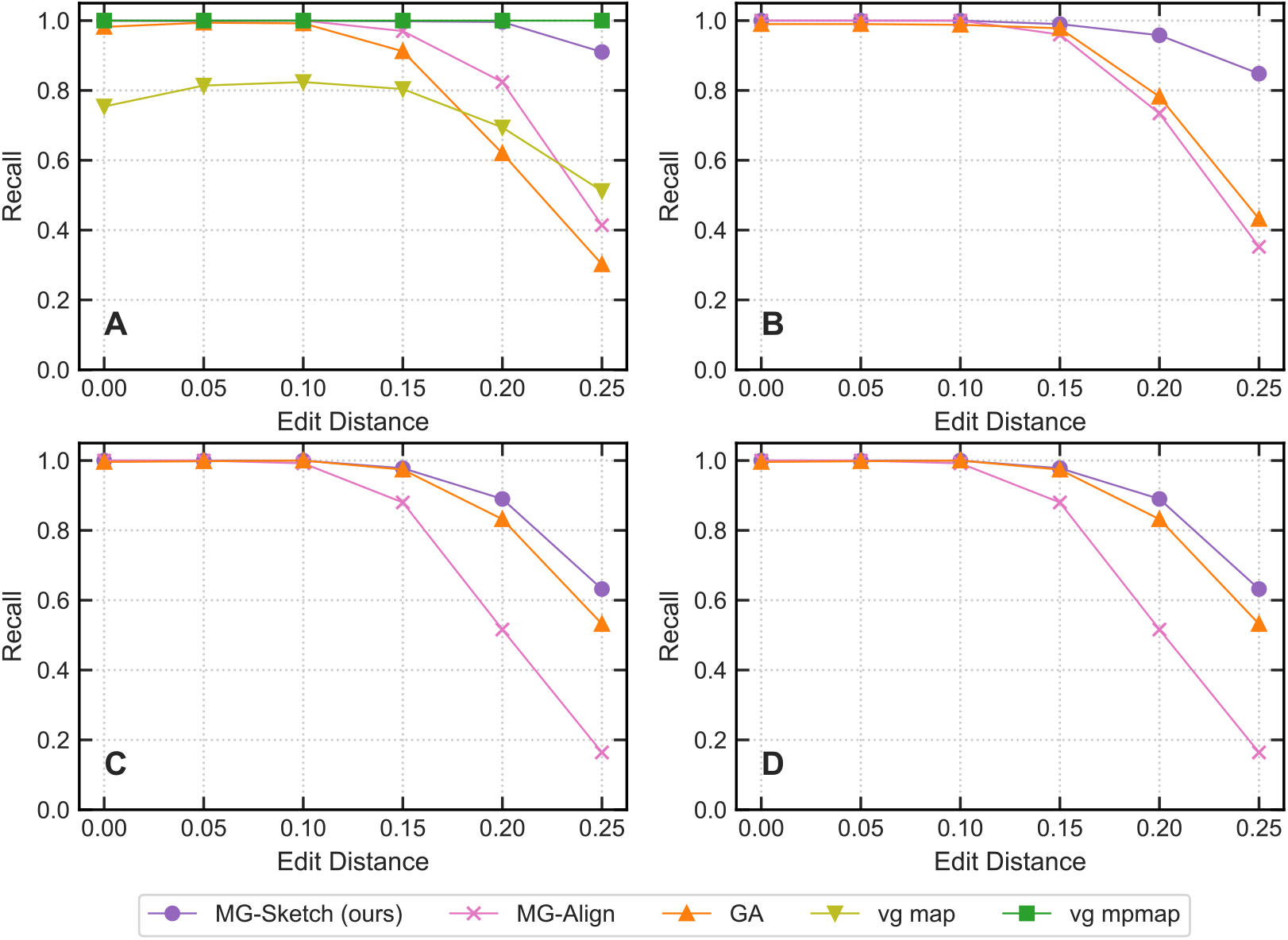
Recall achieved across different mutation rates with increasing graph sizes for each baseline. The number of nodes in the graph for each plot is 100K (A), 10M (B), 100M (C) and 1B (D). Values are measured on the De Bruijn graph generated by MetaGraph. We run MG-Sketch with *K* = 40 neighbors and *D* = 14, *w* = 16, *s* = 8, *t* = 6. Query generation follows the same approach as explained in Section 3.1.

**Fig. 5:**
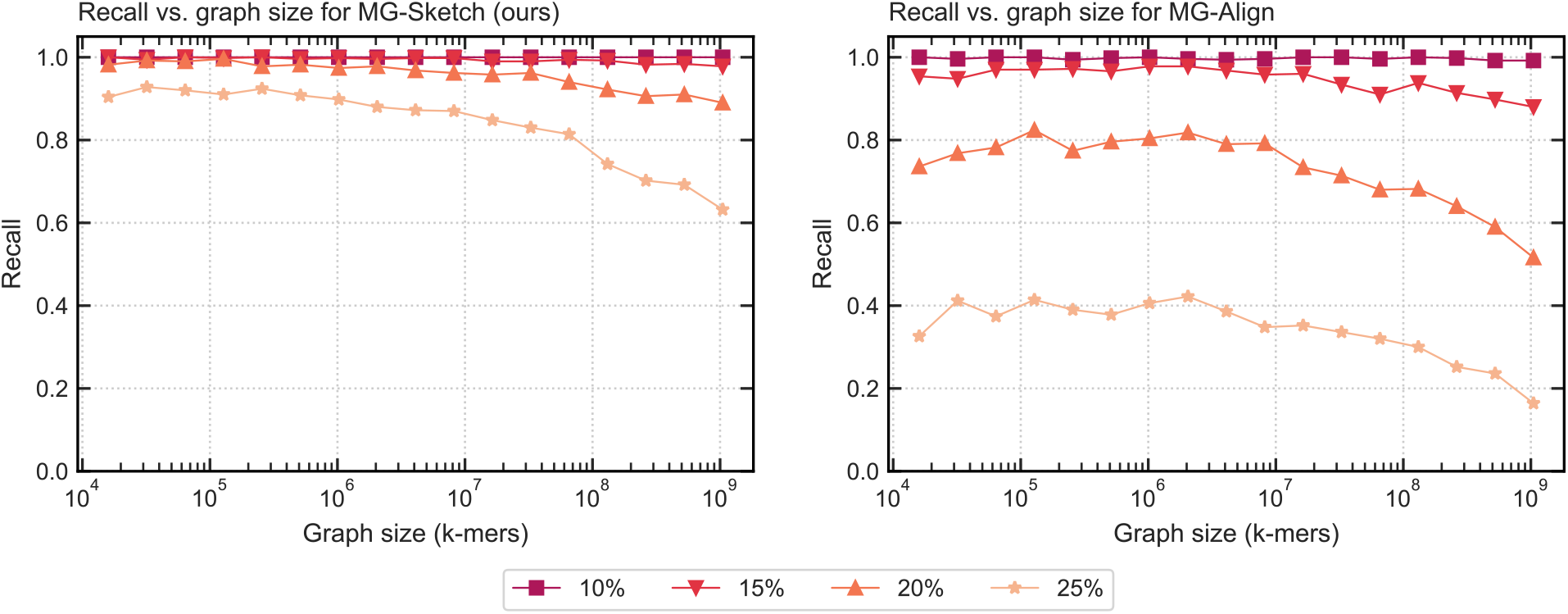
Recall achieved across different mutation rates with increasing graph sizes for MG-Sketch (left) and MG-Align (right). We run MG-Sketch with *t* = 6, *w* = 16, *s* = 8, *D* = 14, and *K* = 40 neighbors. Query generation follows the same approach as explained in Section 3.1. We omit 0% and 5% mutation rate, as both methods achieve an almost perfect recall.

### 3.3 Sketch-based aligner scales quasi-linearly with graph size

We show that the high accuracy of MG-Sketch does not compromise scalability. To compare the scalability of our approach, we measured peak memory usage and average alignment time per query for graphs with 𝓁 ∈ {4,…, 20}, as outlined in Section 3.1. This generates graphs with 16000 up to approximately 1B *k*-mers, posing a challenge to the scalability of each method. We use the MetaGraph base graph for evaluating MG-Sketch and MG-Align (See Appendix A for more details).

#### Peak memory usage scales linearly with graph size

Peak memory usage imposes a hard limit on the size of graphs that a method can preprocess. In Figure 6, we observe that peak memory usage in MG-Sketch scales linearly with the graph size. In particular, for graphs with over 10M nodes, MG-Sketch requires less memory than all other methods except MG-Align. For MG-Align, lower memory usage comes at the cost of lower recall, when compared to MG-Sketch. In contrast, vg mpmap peak memory usage grows rapidly above 80GB. Therefore, we only evaluate vg mpmap for graphs with up to 1M nodes. While vg map requires less memory than vg mpmap, the runtime exceeded 24 hours for graphs with over 60M nodes. It is evident in Figure 6 that the peak memory usage of GA grows faster than linear, making MG-Sketch and MG-Align the only methods with linear memory complexity.

**Fig. 6:**
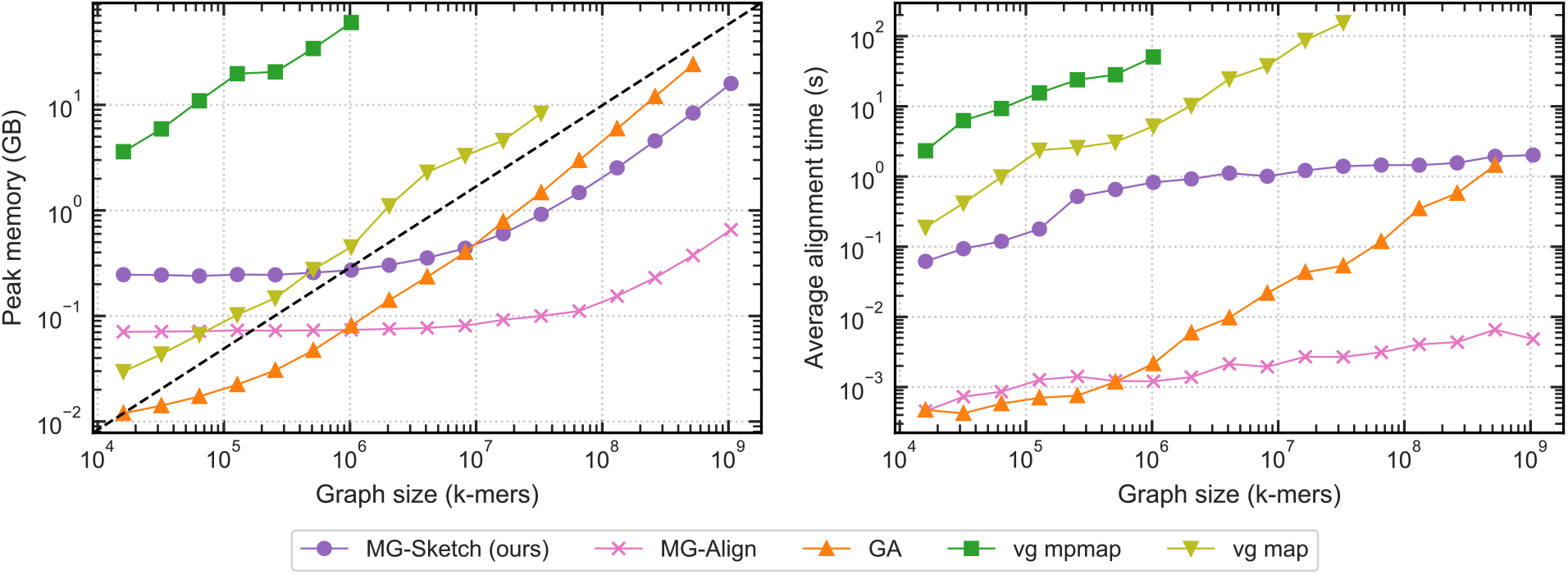
Peak RAM usage and query time vs. graph size. Graphs are generated with *k* = 80 according to Section 3.1. We run MG-Sketch with *t* = 6, *D* = 14, *w* = 16, *s* =8 and *K* = 10 neighbors. Traces for vg mpmap and vg map are incomplete as they exceed time or memory limit. (left) Peak RAM usage vs. graph size. (right) Average alignment time vs. graph size. For recall comparisons see Figure 4.

#### Query alignment time grows logarithmically with graph size

Given that sequence graphs have billions of nodes, it is crucial for the alignment time to grow logarithmically with the graph size. Figure 6 demonstrates that MG-Sketch achieves a logarithmic scale with the graph size. In contrast, the average time for vg map and vg mpmap grows faster than logarithmically. The alignment time for GA also grows faster than linearly, however, this includes the preprocessing time for building the minimizer seeder from the graph.

Notably, the scalability of MG-Sketch does not compromise the recall. Figure 4 demonstrates that for all graph sizes and mutation rates, MG-Sketch achieves a higher or equal recall compared to the other tools, with the exception of vg mpmap. While vg mpmap has higher recall at 25%, for graphs with over 1M it exceeds our memory limit of 80GB.

### 3.4 Good scalability and recall translate into real-world applications

To demonstrate our approach in a more practical setting, we generated a pan-genome De Bruijn graph containing 51,283 assembled virus genome sequences collected from GenBank [35]. Graph construction was performed using the *MetaGraph* framework, utilizing a *k* of 80. The resulting graph contained 337,480,265 nodes. Similar to previous evaluations, we generated 500 query sequences by sampling random walks of length 330 from the graph, subsequently applying random mutations at rates ranging from 5 to 25 percent. Confirming our previous observations on scalability, the sketching-based approach was able to outperform both the MG-Align as well as the GA approaches, starting from mutation rates of 10% and 15%, respectively (Figure 7). Notably, for the highest mutation rate, MG-Sketch almost doubles the recall when compared to the next best approach.

**Fig. 7:**
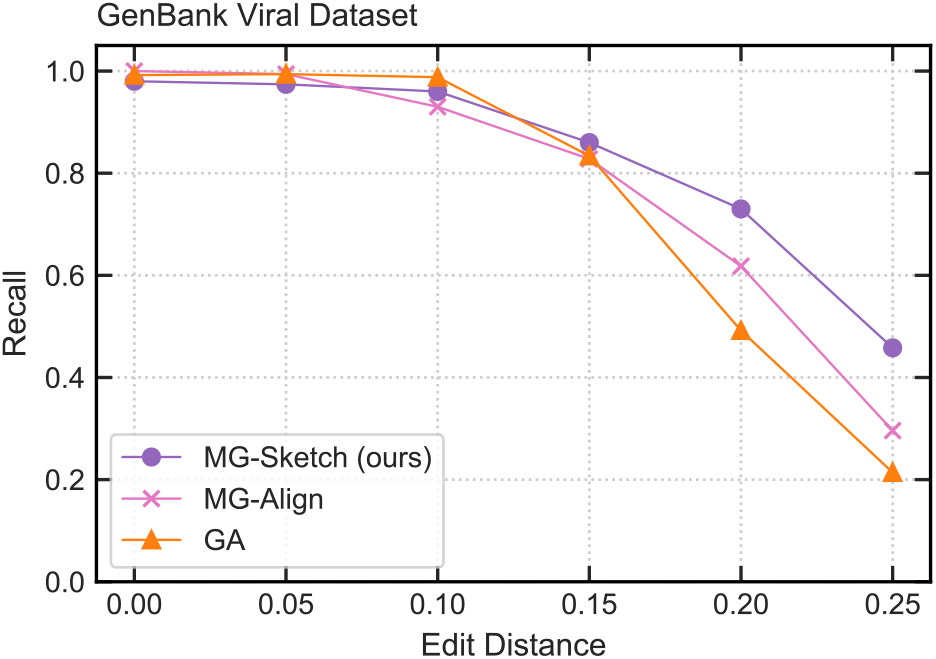
Seeding recall for the GenBank viral graph. Recall is shown for MG-Sketch, MG-Align, and GA as purple, pink, and orange lines, respectively, for query sets of increasing distance, simulated with a random mutation rate ranging from 0% to 25%.

## 4 Conclusion

This work’s main contribution is the introduction of a sketch-based long seed-finding approach that is robust to mutations. We provide a theoretical analysis and demonstrate empirically that our method scales to graphs on the order of 10^9^ nodes. While MG-Align and MG-Sketch share the same extension algorithm, the sketch-based seeds improve alignment recall by a substantial margin in high mutation rate (> 15%) regimes.

While sketching into vector spaces has been successfully applied in *compressive sensing* [7,11], to the best of our knowledge, this is the first work to show that such sketching into vector spaces can lead to improvements in accuracy and scale to genomics-sized datasets. With vector operations, there is a higher gain to be made on systems with specialized hardware, particularly GPUs. In fact, our CUDA-optimized Tensor Sketching implementation is over 200× faster than the single-threaded CPU implementation [17]. Therefore, by employing the CUDA-optimized implementations of FAISS and Tensor Sketching, we can benefit from massive data parallelism on GPUs.

In stark contrast to state-of-the-art tools, our method’s distinguishing feature is that it is a randomized algorithm. At the heart of MG-Sketch is a randomized sketching scheme, which upon running several times independently can boost the recall of the method. This property enables us to prove theoretical guarantees in Lemma 1, despite the method’s conceptual simplicity. Further exploration on the guarantees can bring new insights, such as a probabilistic bound on the deviations for Tensor Sketching, as well as covering indels in the analysis.

There are several aspects that can be improved, but we considered them out the scope of this work. Notably, we have not applied any seed filtering approaches, such as co-linear chaining [32,8,25,1,29]. Furthermore, other sub-sampling strategies for indexing, such as spaced-minimized sub-sampling [14,38], can improve alignment time per query. In conclusion, MG-Sketch takes a novel approach to alignment with promising empirical and theoretical properties. It is also conceptually simple and modular, such that it can be utilized in many other tools. Therefore, it opens up many new interesting areas of research for tackling computational challenges in bioinformatics.

## Acknowledgements

A. J. is funded through Swiss National Science Foundation Project Grant #200550 to A. K. H. M. is funded as part of Swiss National Research Programme (NRP) 75 “Big Data” by the SNSF grant #407540_167331. A. J., H. M., and A. K. are also partially funded by ETH core funding (to G. R.). R.G. was funded through an ETH Research Grant (# ETH-17 21-1) to G.R.

We declare no conflicts of interest.

