## Supplemental Material for "Aligning Distant Sequences to Graphs using Long Seed Sketches"

### Appendix A Commands used for experiments

We use an assembled version of the base graph to evaluate GraphAligner (GA). For the vg methods, we transform the base graph into a variant graph.

To use the same graph in vg map and vg mpmap we "bluntify" it with the GetBlunted tool to remove overlaps between nodes [13]. Finally, we call `vg autoindex` on the blunted graph to obtain a GCSA index used by vg.

We evaluate GA using unlimited `tangle effort` by enabling the parameter `-C -1`. While in this setting GA uses 13GB of RAM, disabling this parameter resulted in an imperfect recall even at 0% mutation. This is due to the heuristic used in tangled graph regions, which can drop the correct path in case the extender starts exploring a false positive path first. This is a common case in our evaluation since our graphs have many similar paths.

We initially generate the De Bruijn graph in MetaGraph. The graph is then assembled into unitigs and blunted using `get_blunted` to accommodate for vg's format. This removes the overlap between the nodes. The GCSA index required by vg is generated using `vg autoindex`. In practice we ran the following set of commands:

```
metagraph build -k 80 --parallel 20 -o graph.dbg sequence.fa
metagraph assemble --to-gfa --compacted --unitigs
        -o graph.gfa graph.dbg
get_blunted --input_gfa graph.gfa > blunted_graph.gfa
vg autoindex -g blunted_graph.gfa -V 2 -w map
vg convert index.xg -p > index.vg
```

To evaluate each baseline we use the following set of commands:

**GraphAligner**

```
GraphAligner -g blunted_graph.gfa -f input.fa
-a output.gaf -x dbg -C -1
```

**MetaGraph Sketching**

```
metagraph align --seeder sketch --embed-dim 14
--num-neighbours 10 --align-end-bonus 0 -i graph.dbg in.fa
```

**MetaGraph Exact**

```
metagraph align --seeder default --align-min-seed-length 15
--align-xdrop 15 -i graph.dbg input.fa
```

**vg map**

```
vg map -z 1 -o 1 -x index.xg -g index.gcsa -f input.fa --gaf
```

**vg mpmap**

```
vg mpmap -n DNA -F GAF -z 1 -o 1 -x index.xg -g index.gcsa
-f input.fa --gaf
```

To measure peak memory usage, we used the `/usr/bin/time -f %M` command. To evaluate vg map and vg mpmap, we extracted the path spelling of the GAF output using `vg find` and `vg view`. To evaluate GraphAligner, we extracted the obtained path spellings from the input GFA file.

### Appendix B Proof of Lemma 1

**Notation** Given a string $a_1 \dots a_n = a \in \Sigma^n$, we define the *I-index* as $a_I = (a_{i_1}, \dots, a_{i_t})$. We write $[X]$ for the indicator variable of event $X$, which is 1 when $X$ holds and 0 otherwise.

**Lemma 1** Let $a$ be a uniform random sequence of length $n$ in $\Sigma^n$, and for a fixed mutation rate $r \in [0, 1]$ let $b$ be a sequence where $a_i$ is substituted by a new character $b_i \in Unif(\Sigma \backslash a_i)$ with probability $r$ and $b_i = a_i$ otherwise. Then for $n \gg 2t\alpha$:

$$\mathbb{E}_{a,b}[d_{te}(a, b)] = (4/\sigma)^{t-1} \cdot r + O(2t\sigma^{2-t}/n) \cdot r,$$

which for DNA with $\alpha = 4$ and fixed $t$ gives $\mathbb{E}[d_{te}(a, b)] = (1 + O(n^{-1})) \cdot r$.

*Proof.* By definition we have

$$\begin{aligned}2\binom{n}{2t-1} d_{te}(a,b) = \|T_a - T_b\|_2^2 &= \sum_{s\in\Sigma^t}\left(\sum_{I\in\mathcal{I}}[a_I = s] - \sum_{I\in\mathcal{I}}[b_I = s]\right)^2 \\ &= \sum_{s\in\Sigma^t}\sum_{I,J\in\mathcal{I}}\Big([a_I = s][a_J = s] - [a_I = s][b_J = s] - [b_I = s][a_J = s] + [b_I = s][b_J = s]\Big).\end{aligned}$$

By symmetry between $a$ and $b$, the first and last term, and second and third term are equal in expected value, reducing this to

$$\begin{aligned}\mathbb{E}_{a,b}\left(\|T_a - T_b\|_2^2\right) &= \mathbb{E}\left(2\sum_{s\in\Sigma^t}\sum_{I,J\in\mathcal{I}}\Big([a_I = s][a_J = s] - [a_I = s][b_J = s]\Big)\right) \\ &= \mathbb{E}\left(2\sum_{I,J\in\mathcal{I}}\sum_{s\in\Sigma^t}\Big([a_I = s \wedge a_J = s] - [a_I = s \wedge b_J = s]\Big)\right) \\ &= 2\sum_{I,J\in\mathcal{I}}\mathbb{E}\Big([a_I = a_J] - [a_I = b_J]\Big). \qquad (3)\end{aligned}$$

Define the overlap $q$ as the number of positions where $I$ and $J$ are equal, $q(I, J) := |\{x \in [t] : I_x = J_x\}|$. We will show using induction on $t$ that $\mathbb{E}[a_I = b_J] = (\sigma(1 - r))^q\sigma^{-t}$. For $t = 0$ we have $I = J = \varnothing$ and trivially $\mathbb{E}[a_I = b_J] = 1$. For $t > 0$, write $I'$ and $J'$ for the tuples $(I_1, \dots, I_{t-1})$ and $(J_1, \dots, J_{t-1})$. When $I_t = J_t$, the characters $a_{I_t}$ and $b_{J_t}$ are independent of the earlier characters and equal with probability $1 - r$, and $q(I', J') = q - 1$, so that

$$\begin{aligned}\mathbb{E}[a_I = b_J] &= (1 - r)\mathbb{E}[a_{I'} = b_{J'}] \\ &= (1 - r) \cdot (\sigma(1 - r))^{q-1}\sigma^{-(t-1)} \\ &= (\sigma(1 - r))^q\sigma^{-t}.\end{aligned}$$

When $I_t \neq J_t$, assume without loss of generality that $I_t < J_t$. Then $I_x < J_t$ for all $x \in [t]$, resulting in $b_{J_t}$ is independent from the characters seen so far. This implies that $[a_{I_t} = b_{J_t}]$ is independent from $[a_{I'} = b_{I'}]$:

$$\begin{aligned}\mathbb{E}[a_I = b_J] &= \mathbb{E}[a_{I_t} = b_{J_t}]\mathbb{E}[a_{I'} = b_{J'}] \\ &= \sigma \cdot (\sigma(1 - r))^q\sigma^{-(t-1)} \\ &= (\sigma(1 - r))^q\sigma^{-t}.\end{aligned}$$

We conclude that

$$\mathbb{E}_{a,b}\big([a_I = a_J] - [a_I = b_J]\big) = \sigma^{-t+q}\big(1 - (1 - r)^{q(I,J)}\big).$$

This difference vanishes for $q = 0$, and thus in (3) we only have to consider $(I, J)$ with $q(I, J) \geq 1$. The summation can now be rewritten as

$$\begin{aligned}
\mathbb{E}_{a,b}\left(\|T_a - T_b\|_2^2\right) &= 2\sum_{q=1}^{t} \sum_{\substack{I,J\in\mathcal{I}:\\ q(I,J)=q}} \mathbb{E}\Big([a_I = a_J] - [a_I = b_J]\Big) \\
&= 2\sum_{q=1}^{t} \sum_{\substack{I,J\in\mathcal{I}:\\ q(I,J)=q}} \sigma^{-t+q}\big(1-(1-r)^q\big) \\
&= 2\sum_{q=1}^{t} \sigma^{-t+q}\big(1-(1-r)^q\big)\cdot f_q, \qquad (4)
\end{aligned}$$

where $f_q$ counts the number of pairs $(I, J)$ with $q(I, J) = q$. Since $|I \cap J| \geq q$, the total number of distinct indices is bounded by $|I \cup J| \leq 2t - q$. This directly implies that $f_q \leq (1 + o(1))\binom{n}{2t-q}$, which for $q \geq 2$ gives

$$\binom{n}{2t-1}^{-1}\binom{n}{2t-q}\cdot\sigma^{-t+q}\big(1-(1-r)^q\big) = O((2t\sigma/n)^{q-1}\sigma^{1-t}r).$$

When $q = 1$ and $|I \cup J| < 2t - 1$ a similar argument applies, and we are left with the case where $q = 1$ and $|I \cup J| = 2t - 1$. We can first choose the $2t - 1$ distinct values for $I \cup J$ in $\binom{n}{2t-1}$ ways, and then assume that $I \cup J = [2t - 1]$. The overlap can be at any odd position $2k + 1 \in \{1, 3, \ldots, 2t - 1\}$, since $I$ and $J$ must both have an equal number of distinct elements smaller (resp. larger) than $2k + 1$. Given the overlap at $2k + 1$, the $2k$ smaller positions can be split into two halves in $\binom{2k}{k}$ ways, and similarly for the right half, leading to the following number of $(I, J)$ pairs with $q = 1$ and $|I \cup J| = 2t - 1$:

$$\binom{n}{2t-1}\cdot\sum_{k=0}^{t-1}\binom{2k}{k}\binom{2(t-1-k)}{t-1-k} = \binom{n}{2t-1}\cdot 4^{t-1},$$

a well-known identity [21,12]. Splitting (4) into the cases $q = 1$ (with $|I \cap J| = 1$ and $|I \cap J| > 1$) and $q \geq 2$, and assuming that $n \gg 2t\sigma$, we get our result:

$$\begin{aligned}
\mathbb{E}(d_{te}(a,b)) &= (4/\sigma)^{t-1}\cdot r + O(2t\sigma/n\cdot\sigma^{-t}r) + \sum_{q=2}^{t} O((2t\sigma/n)^{q-1}\cdot\sigma^{1-t}r) \\
&= (4/\sigma)^{t-1}\cdot r + O(2t\sigma^{2-t}/n)\cdot r. \qquad \square
\end{aligned}$$

### Appendix C Implementation details for MG-Sketch

We provide an overview of the implementation of Tensor Slide Sketching.

---

**Algorithm 2:** TensorSlideSketch $\phi_{TSS}$

---

**Input** : Query $q \in \Sigma^k$
**Output** : Sketch $\Phi \in \mathbb{R}^{\lceil \frac{k-w+1}{s} \rceil D}$
**Parameter:** Dimension $D \in \mathbb{N}^+$
Tuple size $t \in \mathbb{N}^+$
Stride $s \in \mathbb{N}^+$
Window size $w \in \mathbb{N}^+$
**for** $j \leftarrow 0$ *to* $\lceil \frac{k-w+1}{s} \rceil$ **do**
 | $\Phi_j \leftarrow$ TensorSketch$\big(q[js : js + w - 1]\big)$
**end**
**Return** : $\Phi$

---
